# Full-length transcriptome atlas of *Panax vietnamensis* var. *fuscidiscus* reveals novel genes and alternative splicing in tissue-specific biosynthesis of ocotillol-type saponins

**DOI:** 10.64898/2026.01.19.700240

**Authors:** Wei Li, Zhe Xu, Aneela Nijabat, Li-zhi Gao

## Abstract

**Background:** *Panax vietnamensis* var. *fuscidiscus* is the only known *Panax* species that accumulates exceptionally high levels of ocotillol (OCT)-type ginsenosides, particularly majonoside R2. However, the molecular mechanisms underlying OCT-type ginsenoside biosynthesis remain largely uncharacterized, due in part to the absence of comprehensive full-length transcriptome data and complete gene annotations.

**Results:** We combined PacBio single-molecule real-time (SMRT) and Illumina RNA sequencing (RNA-Seq) technologies to construct the first tissue-specific, full-length transcriptome atlas of *P. vietnamensis* var. *fuscidiscus*. A total of 281,468 non-redundant transcripts were assembled, including 8,089 novel genes not present in existing annotations, substantially expanding the documented gene repertoire of this species. We identified 21 candidate genes putatively involved in triterpenoid saponin biosynthesis and uncovered distinct tissue-specific expression patterns across the oleanane (OA)-, protopanaxadiol (PPD)-, protopanaxatriol (PPT)-, and OCT-type biosynthetic pathways. Notably, we detected 46,917 alternative splicing (AS) events, with over 45% of expressed genes producing multiple isoforms. Several key biosynthetic genes—including dammarenediol synthase (*DDS*), cytochrome P450 716A53v2 (*CYP716A53v2*), and mevalonate diphosphate decarboxylase (*MVD*)—exhibited tissue-specific splicing patterns, such as retained introns and exon skipping, which may contribute to functional diversification of isoforms involved in OCT-type ginsenoside biosynthesis. These results highlight the importance of post-transcriptional regulation and transcript diversity in the evolution and fine-tuning of specialized metabolic pathways.

**Conclusions:** This study establishes a high-quality, full-length transcriptomic resource for *P. vietnamensis* var. *fuscidiscus*, revealing previously undescribed genes and extensive alternative splicing events related to ginsenoside biosynthesis. Our findings provide novel molecular insights into the mechanisms regulating OCT-type saponin accumulation and offer a foundation for future research into the functional characterization, regulatory mechanisms, and synthetic biological production of rare triterpenoid saponins.

## Background

Medicinal plants play a crucial role in promoting human health. In developing countries, herbal medicines serve as the primary form of healthcare for approximately 75% of the population. Known as the “King of Herbs”, ginseng (*Panax* spp.) has been used in traditional Chinese medicine for over 5,000 years and has gained increasing acceptance in Western medicine in recent decades. The main bioactive compounds in ginseng are ginsenosides, a group of triterpenoid saponins that are predominantly found in *Panax* species, with the highest concentrations typically located in the roots and rhizomes [1, 2]. To date, more than 300 natural ginsenosides have been identified across *Panax* species [3–5]. Based on their aglycone backbone structures, ginsenosides are categorized into two major types: dammarane-type and oleanane (OA)-type. The dammarane-type can be further divided into three subclasses: protopanaxadiol (PPD)-, protopanaxatriol (PPT)-, and ocotillol (OCT)-type ginsenosides [3].

In *Panax*, OCT-type ginsenosides represent a specialized subclass of dammarane-type ginsenosides. Oxidosqualene cyclases (*OSCs*), which initiate dammarane-type ginsenoside biosynthesis, have been shown to generate OCT-related precursors through functional diversification following gene duplication. However, the downstream biosynthetic pathways and regulatory mechanisms governing OCT-type ginsenoside accumulation remain poorly understood [6, 7].

*Panax vietnamensis* Ha et Grushv., a new *Panax* species, was first discovered in the central highlands of Vietnam in 1973 and is the southernmost distributed species within the genus [8]. The roots and rhizomes of this species have long been secretly used by the Sedang ethnic group in Vietnam as a life-saving medicinal plant, and it holds significant ethnomedical value [9–11]. Notably, *P. vietnamensis* is the only known *Panax* species that accumulates high levels of OCT-type ginsenosides [12]. Among them, the representative compound majonoside R2 accounts for over 5% of the dry weight of the rhizome and approximately half of the total ginsenosides, making *P. vietnamensis* an important resource for studying OCT-type ginsenosides [13–15].

Interestingly, a new variety of *P. vietnamensis*, named *P. vietnamensis* var. *fuscidiscus* K. Komatsu, S. Zhu & S.Q. Cai (commonly known as “Ye Sanqi” in Chinese), was discovered in Jinping County, Southern Yunnan Province, China [16]. However, subsequent studies suggest that its taxonomic classification as a variety of *P. vietnamensis* requires further investigation [17]. This variety contains significantly higher levels of majonoside R2 compared to other genotypes, making it an ideal model for studying the biosynthetic mechanisms of OCT-type triterpene saponins [18, 19]. Although recent studies have identified two UDP-glycosyltransferases (*PvfUGT1* and *PvfUGT2*) responsible for the sequential glycosylation steps leading to majonoside R2 formation, the upstream regulatory architecture of this pathway remains largely unresolved. OCT-type saponins were first identified as minor constituents from the underground parts of *P. japonicus* var. *major* [20]. Pharmacological studies have since demonstrated their multiple bioactivities, including anti-tumor effects [21], anti-stress properties [22], and hepatoprotective functions [23, 24].

Although substantial research has been focused on the diversity, content, metabolic pathways, and pharmacological activities of ginsenosides in widely used species, such as *P. ginseng*, *P. quinquefolius*, and *P. notoginseng* [3, 25–28], studies on the biosynthetic pathways of OCT-type ginsenosides in *P. vietnamensis* remain limited. Recent studies have applied the Illumina sequencing platform to *P. vietnamensis*, including the release of draft genome assembly [19, 29], tissue-specific transcriptome analyses, and mining of genes involved in ginsenoside biosynthesis [19, 30, 31]. However, the limitations of short-read sequencing, such as its inability to fully capture complex splicing isoforms and full-length transcripts, have hindered the comprehensive understanding of the molecular mechanisms underlying OCT-type ginsenoside biosynthesis in *P. vietnamensis*.

Single-molecule real-time (SMRT) sequencing, a third-generation sequencing technology, enables the direct acquisition of full-length cDNA transcripts and overcomes the shortcomings of short-read sequencing in transcript assembly and isoform identification. In recent years, SMRT sequencing has been successfully applied in various plant research areas, including gene structure analysis, alternative splicing (AS), transcription factor identification, and simple sequence repeat (SSR) discovery [32–35]. In this study, we report the first integration of SMRT and second-generation sequencing (SGS) technologies to generate a high-quality, full-length, tissue-resolved transcriptome of *P. vietnamensis* var. *fuscidiscus*. By combining PacBio Isoform Sequencing (Iso-Seq) for isoform-level transcript discovery and Illumina RNA sequencing (RNA-Seq) for expression quantification and annotation, we constructed a comprehensive transcriptomic atlas across the five major tissues (root, rhizome, stem, leaf, and flower). This integrative approach enabled the identification of 8,089 novel genes and over 46,000 alternative splicing events, significantly improving the resolution of gene structure and regulatory complexity. Importantly, we identified key candidate genes and splicing isoforms potentially involved in the biosynthesis of OCT-type triterpenoid saponins, offering valuable resources for functional genomics and metabolic engineering in *Panax* species.

## Methods

### Plant materials

Three-year-old *P. vietnamensis* var. *fuscidiscus* plants were collected in October 2015 from the cultivation base of Jinping Shili Medicinal Material Development Co., Ltd. in Jingping County, Yunnan Province, China (22°44′24″ N, 103°13′48″ E; Altitude: 1750 m). The plants were grown under the forest canopy in humus-rich soil, with approximately 25 - 30 cm spacing between individuals, and maintained under natural temperature and light conditions typical of the local environment. During the sampling period, the average temperature was 16 - 24 ℃ with a photoperiod of about 12 hours of daylight. Root, stem, leaf, flower, and rhizome tissues were cut into small pieces, immediately frozen in liquid nitrogen, and stored at − 80 ℃ until RNA extraction.

### Construction of Illumina RNA-Seq library

Total RNA was extracted from frozen root, stem, leaf, flower, and rhizome tissues using the Plant RNA Extraction Kit (Zoonbio Biotechnology, Cat. No. RK16-50T) following the manufacturer’s instructions. The integrity, quality, and concentration of the extracted RNA were assessed using the Agilent 2100 Bioanalyzer system (Agilent Technologies, Santa Clara, CA, USA). High-quality RNA samples were used to construct cDNA libraries with the Illumina mRNA-Seq Sample Preparation Kit (Illumina, San Diego, CA, USA). Sequencing was performed on the Illumina HiSeq 2000 platform using standard protocols.

Raw reads were processed with Trimmomatic (v0.39) to remove adapter sequences, trim low-quality bases (Phred score < 15), and cut the first 15 bases from each read (HEADCROP:15). Cleaned reads were *de novo* assembled using Trinity (v2.15.2), and redundant sequences were removed using CD-HIT (v4.8.1) [36].

### cDNA library preparation and PacBio SMRT sequencing

Total RNA was extracted using TRIzol Reagent (Invitrogen, Beijing, China; Cat. No. 15596-026) according to the manufacturer’s instructions. RNA integrity was evaluated with an Agilent 2200 TapeStation (Agilent Technologies, USA). Full-length cDNA libraries were prepared following the PacBio Isoform Sequencing (Iso-Seq) protocol, using the Clontech SMARTer PCR cDNA Synthesis Kit (Cat. No. 634925) and the BluePippin Size Selection System (Sage Science, USA), with minor modifications. Equal amounts of high-quality total RNA (10 μg, RNA Integrity Number > 8.5) were pooled from four tissues (root, stem, leaf, and flower). The cDNA was size-selected into three fractions: 1-2 kb, 2-3 kb, and 3-6 kb using BluePippin. Each size fraction (500 ng) was used to construct SMRTbell libraries with the PacBio Template Prep Kit 1.0 (Part No. 100-259-100). A total of 13 SMRT cells were sequenced on the PacBio RS II platform: five cells for both 1–2 kb and 2–3 kb libraries, and three cells for the 3-6 kb library. Sequencing was performed using the P4-C2 chemistry (BINDINGKIT=100356300; SEQUENCINGKIT=100356200) with 120-minute movies on the PacBio RS II system. An additional eight SMRT cells were run: three for each of the 1-2 kb and 2-3 kb libraries, and two for the 3-6 kb library. Raw reads were processed using the SMRT Portal analysis suite (PacBio) to generate subreads for downstream analysis.

### Data analysis of PacBio SMRT long reads

PacBio single-molecule long reads were analyzed using RS_IsoSeq (v2.3) and the smrtanalysis_2.3.0.140936.p4.150482 pipeline. The script pbtranscript.py was used for the characterization of full-length (FL) reads, with 5′/3′ primers and poly(A) tails employed to discriminate strand-specific full-length reads.

Circular consensus sequencing (CCS) was executed by SMRT Link, and high-quality, FL, and polished consensus transcripts were obtained using the isoform-level clustering algorithm ICE (Iterative Clustering for Error Correction). FL non-chimeric (FLNC) reads were identified, and high-quality (HQ) and low-quality (LQ) consensus isoforms were generated after clustering. LQ isoforms were further corrected with cleaned SGS reads using LoRDEC (v0.9) [37]. HQ and corrected LQ isoforms were collapsed by CD-HIT (v4.8.1) to form the final PacBio isoforms.

### Merging RNA-Seq assembly and Iso-Seq isoforms

The assembled transcripts from second-generation sequencing (SGS) data and collapsed PacBio isoforms were combined. CD-HIT (v4.8.1) was then used to cluster the transcripts and to remove redundancies. The merged transcript set was classified into coding and non-coding RNAs using RNASamba (v0.2.4, classify mode) with a pre-trained full-length model (full_length_weights.hdf5). Coding sequences were exported as predicted proteins in FASTA format. This non-redundant, merged transcript set was subsequently used as the reference for downstream gene expression analysis.

### Gene functional annotation and expression analysis

Gene functions were annotated using eggNOG-mapper (v2.1.12) with DIAMOND mode [38], based on a locally installed eggNOG database. The annotation output was processed with the emapperx.R script provided in the emcp package (http://git.genek.cn:3333/zhxd2/emcp), which generated Gene Ontology (GO) assignments, Kyoto Encyclopedia of Genes and Genomes (KEGG) pathway annotations, and a species-specific OrgDB database for downstream enrichment analyses. Expression quantification across tissues was performed using RSEM (v1.3.1) with SGS reads [39].

### KEGG functional classification of expressed genes

To characterize the functional profiles of different tissues, KEGG pathway classification was performed on genes that were expressed (TPM > 2) in at least one of the five tissues. KEGG annotation was conducted using TBtools-II (v2.376) with the Plants.20211128.TBtoolsKeggBackEnd database to ensure plant-specific pathway assignment [40]. KO annotations from eggNOG-mapper were mapped to KEGG Level 1 and Level 2 categories through the plant-specific backend, and non-plant categories were automatically excluded. The resulting KEGG functional distribution was used to compare pathway composition among tissues.

### Differential expression analysis and KEGG enrichment analysis

Differential expression analysis between tissues was performed using edgeR [41], implemented through the Trinity utility script run_DE_analysis.pl with a fixed dispersion value of 0.1. Genes with |log₂FC| ≥ 1 and false discovery rate (FDR) ≤ 0.05 were defined as differentially expressed genes (DEGs), while others were considered not significant. As only one sample per tissue was sequenced, no biological replicates were included.

KEGG enrichment analysis for DEGs was performed using the enricher () function in the clusterProfiler package with the customized OrgDB database generated by emcp, ensuring accurate mapping to plant KEGG orthology. For enrichment analysis, the DEGs list was used as the input gene set, and all annotated genes were supplied as the background (universe). Significance thresholds were set to pvalueCutoff = 0.05 and qvalueCutoff = 0.05, and enrichment significance was further adjusted using the Benjamini–Hochberg method.

### Full-length transcript reconstruction and isoform identification

Full-length transcript sequences were aligned to the *P. vietnamensis* var. *Fuscidiscus* reference genome (https://db.cngb.org/data_resources/project/CNP0002878/) [17] using minimap2 (v2.28-r1209) with spliced alignment parameters (-ax splice -uf -k14 --secondary=no) [42]. Resulting SAM files were converted to sorted and indexed BAM format using SAMtools (v1.21) [43]. Redundant isoforms were collapsed using Cupcake’s collapse_isoforms_by_sam.py (https://github.com/Magdoll/cDNA_Cupcake), generating high-confidence non-redundant transcript loci.

The collapsed isoforms were processed with Cogent to cluster transcripts into gene families and reconstruct representative gene models. Specifically, run_mash.py computed pairwise k-mer distances (k=30), and process_kmer_to_graph.py transformed these distances into similarity graphs. Connected components within the graph were reconstructed using reconstruct_contig.py.

To evaluate isoform structure and classify transcript types, SQANTI3 [35] was employed with the reference genome and GTF annotation as input. SGS reads were used to support splice junction validation and saturation analysis. Low-confidence isoforms were removed based on SQANTI3’s filtering rules, resulting in a high-confidence, structurally annotated transcriptome.

### Identification of alternative splicing events

Alternative splicing (AS) events were identified using SUPPA2 [44], based on the SQANTI3-corrected transcript annotations in GTF format. The generateEvents function was used to detect and categorize seven major AS types: skipped exon (SE), retained intron (RI), mutually exclusive exons (MX), alternative 3′ splice site (A3), alternative 5′ splice site (A5), alternative first exon (AF), and alternative last exon (AL).

### Identification of saponin biosynthesis-related genes

Candidate genes potentially involved in saponin biosynthesis were identified through homology-based searches using BLASTP (v2.16.0). A local protein database was first constructed from the full set of predicted transcript-derived protein sequences using the makeblastdb tool. Fifty-six known saponin biosynthetic protein sequences were used as queries against this database with an E-value threshold of 1e-5.

Putative oxidosqualene cyclases (*OSCs*) were further identified using HMMER (v3.3.2) [45]. Protein sequences were scanned against the squalene-hopene cyclase N-terminal (PF13249) and C-terminal (PF13243) domain models obtained from the Pfam database (v37.0) [46, 47] with the trusted cutoff parameter (-cut_tc). Only sequences containing both domains were considered high-confidence *OSCs* and retained for downstream analysis.

Putative UDP-glycosyltransferases (*UGTs*) were identified using HMMER (v3.3.2). All predicted protein sequences were scanned against the PSPG-domain HMM model (PF00201) downloaded from the Pfam database (v37.0) using the trusted cutoff parameter (-cut_tc). Hits were further filtered by significance (E-value < 1×10^-20^) and domain length (350-600 aa) to remove truncated or non-canonical sequences. To ensure glycosyltransferase specificity, candidate proteins were additionally screened for the conserved PSPG-box motif (HCGWNSX_0-3_) using regular-expression matching. Expression data (TPM) from the five tissues were then incorporated, and only genes with TPM > 2 in at least one tissue were retained as high-confidence *UGTs*.

The final *UGTs* set was subjected to multiple sequence alignment using MAFFT (-auto) and phylogenetic reconstruction using IQ-TREE (-m MFP -bb 1000) for downstream classification.Visualization of the phylogenetic tree and gene expression heatmap was performed using tvBOT (v2.6.1) [48].

### Statistical analysis

All statistical analyses and visualizations were performed in R (v4.5.0) using the packages tidyverse [49], ggplot2 [50], EnhancedVolcano [51], pheatmap [52], and clusterProfiler [53].

## Results

### Full-length transcriptome sequencing and integrated assembly

To construct a comprehensive transcriptome of *P. vietnamensis* var. *fuscidicus*, we employed a hybrid sequencing strategy integrating third-generation PacBio single-molecule real-time (SMRT) sequencing and second-generation sequencing (SGS). For SMRT sequencing, high-quality total RNA was extracted from root, stem, leaf, and flower tissues of three-year-old plants at the flowering stage. Equal amounts of RNA from each tissue were pooled to generate three size-fractionated cDNA libraries (1–2 kb, 2–3 kb, and 3–6 kb). These libraries were sequenced on the PacBio RS II platform using 13 SMRT cells, producing 619,574 reads of insert (ROIs), including 310,550 full-length non-chimeric (FLNC) reads identified by the presence of both 5′ and 3′ primers and a poly(A) tail (**Table S1**). The length distribution of FLNC reads corresponded well with the expected size ranges of the respective libraries (**Fig. 1A**, **1B**), confirming effective size selection. An additional 282,547 non-full-length reads were also obtained. After iterative clustering and error correction (ICE) and LoRDEC polishing with Illumina reads, 109,990 high-confidence isoforms were retained, with lengths ranging from 180 to 19,929 bp (**Table S1**; **Fig. 1A**).

**Figure 1.**
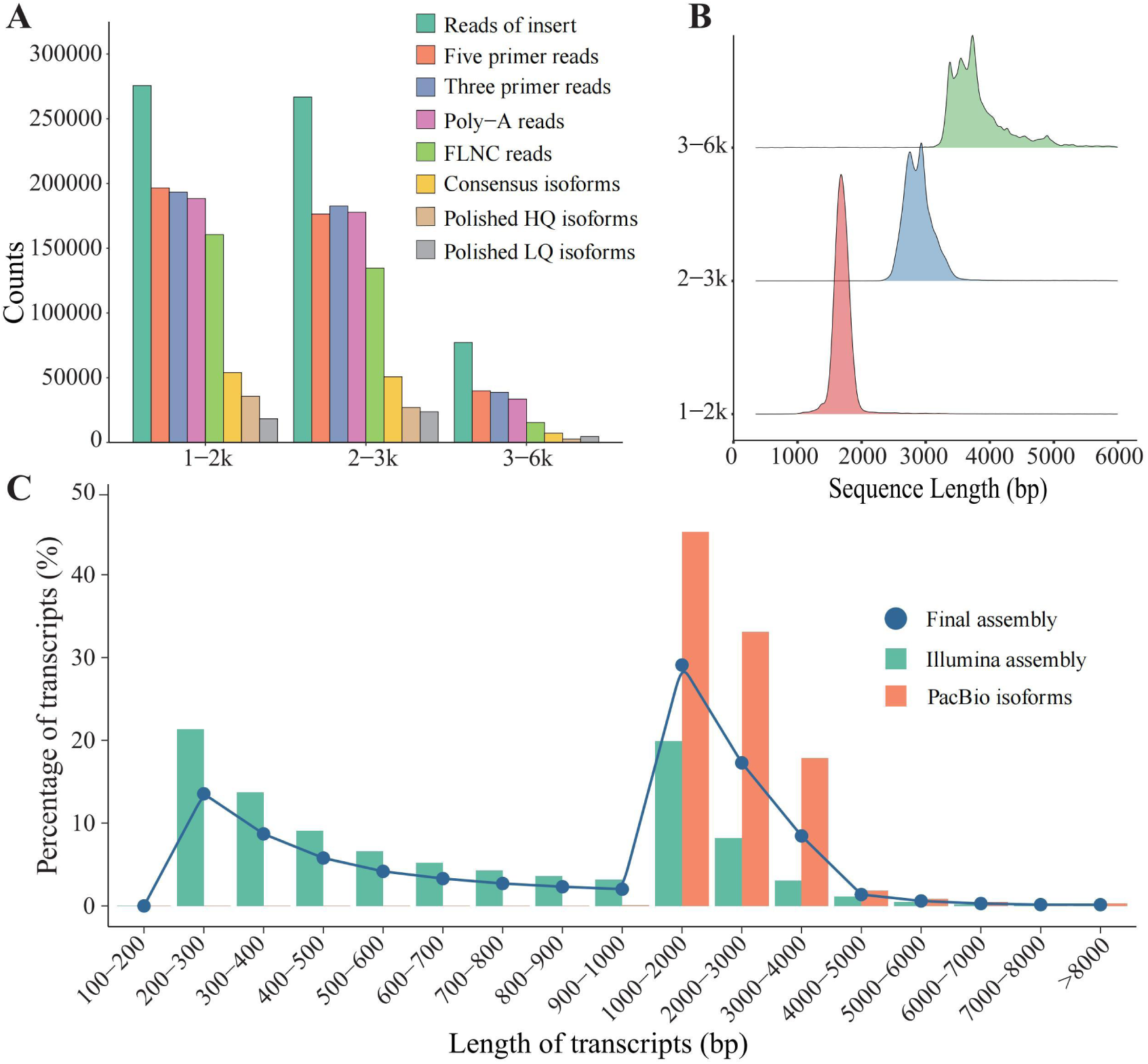
Overview of PacBio SMRT sequencing and transcript length distribution. (A) Distributions of read counts and lengths from PacBio libraries with size fractions of 1–2 kb, 2–3 kb, and 3–6 kb. (B) Length distribution of full-length non-chimeric (FLNC) reads across different libraries. (C) Comparison of transcript lengths between PacBio SMRT and Illumina assemblies.

For SGS, total RNA from five tissues (root, stem, leaf, flower, and rhizome) was sequenced individually on the Illumina platform, yielding 97,206,112 cleaned reads. *De novo* assembly using Trinity produced 195,406 contigs with a total length of approximately 197 Mb. Redundancy reduction with CD-HIT resulted in a final set of 191,832 unigenes (**Table S1**).

Merging PacBio isoforms and Illumina-assembled unigenes, followed by clustering with CD-HIT, yielded a final reference transcriptome comprising 281,468 non-redundant transcripts with a total length of ∼410 Mb, a GC content of 40.06%, and an N50 of 2,471 bp (**Table S1**). The merged transcriptome showed a broader and more uniform transcript length distribution than the individual assemblies, effectively balancing the short contigs from Illumina and the long isoforms from PacBio (**Fig. 1C**).

### Reconstruction and annotation of transcripts

Stringent filtering with SQANTI3 yielded a high-confidence reference transcriptome comprising 70,835 isoforms derived from 26,258 genes. Notably, 8,089 genes were newly identified and absent from the reference annotation, suggesting that Iso-Seq is effective in facilitating the detection of previously unannotated genomic loci.

Analysis of isoform counts showed that 52.0% of genes produced a single isoform, while 48.0% generated two or more isoforms (ranging up to 16), indicating a high prevalence of alternative splicing (AS) and a complex transcriptional landscape in *P. vietnamensis* (**Fig. 2A**). SQANTI3 identified 120,846 splice junctions from the Iso-Seq data. Among these, 67.4% (81,448) were previously annotated as canonical (GT–AG) junctions, 32.5% (39,274) were novel canonical junctions, and only 0.1% (124) were novel non-canonical junctions (**Fig. 2B**). Isoform classification revealed that 46.43% were novel transcripts, including 9.39% from known genes (Novel In Catalog, NIC) and 37.04% from unannotated loci (Novel Not in Catalog, NNC), underscoring the substantial novelty captured by Iso-Seq. Known isoforms accounted for 36.78% of the dataset and fell into the full splice match (FSM) and incomplete splice match (ISM) categories, while the remaining transcripts were classified as non-canonical types, including antisense, fusion, and intergenic transcripts (**Fig. 2C**).

**Figure 2.**
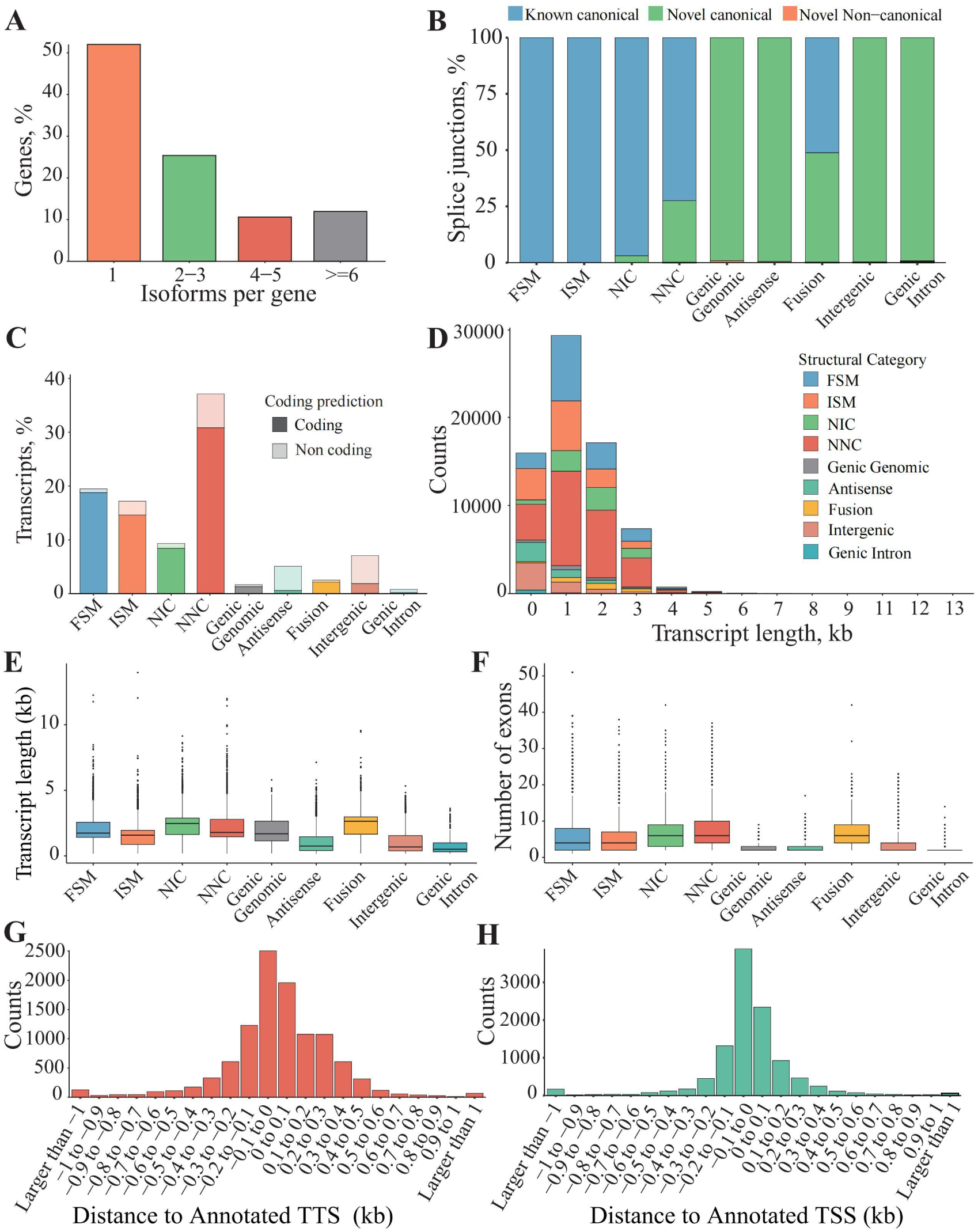
Structural characterization of Iso-Seq isoforms in *P. vietnamensis* var. *fuscidiscus*. (A) Gene-level splicing complexity, represented by the number of isoforms per gene. (B) Comparison of isoform counts between annotated and novel genes. (C) SQANTI3 classification of isoforms into nine structural categories: full-splice match (FSM), incomplete splice match (ISM), novel in catalog (NIC), novel not in catalog (NNC), genic genomic, antisense, fusion, intergenic, and intronic transcripts. Each category is subdivided into protein-coding (dark) and non-coding (light) isoforms. (D) Isoform length distribution across SQANTI3 categories. (E) Summary of isoform length ranges by structural class. (F) Exon count distribution among isoforms across categories. (G) Distance between 3′ ends of FSM transcripts and annotated transcription termination sites (TTS); negative values indicate upstream positions. (H) Distance between 5′ ends of FSM transcripts and annotated transcription start sites (TSS); negative values indicate downstream positions relative to annotated TSS.

Further analysis showed that most isoforms were 1–3 kb, with FSM and ISM generally shorter. NIC and NNC isoforms showed a higher proportion of long transcripts, as indicated by the extended upper tails in transcript length distribution (Fig. 2E). Although their exon number distributions largely overlapped with those of FSM and ISM (Fig. 2F), NIC and NNC still included numerous multi-exon isoforms, suggesting a tendency toward increased structural complexity. This highlights long-read sequencing’s ability to capture structurally complex isoforms beyond the predominant size range (Fig. 2D–F). Transcript boundary analysis revealed that FSM isoforms had 3′ ends that closely matched annotated transcription termination sites (TTS), supporting a high level of accuracy in termination detection. In comparison, their 5′ ends often extended upstream of annotated transcription start sites (TSS), suggesting that Iso-Seq may provide more complete transcript boundary information (**Fig. 2G**, **2H**).

Together, these results highlight the value of long-read Iso-Seq in capturing full-length transcripts, discovering novel isoforms, and elucidating the complex transcriptomic architecture of *P. vietnamensis*.

### Functional characterization and tissue-specific transcript profiles

A total of 131,784 transcripts (80.55%) were annotated in EggNOG or COG/KOG databases, 84,826 (51.85%) were assigned Gene Ontology (GO) terms, and 52,683 (32.20%) were mapped to KEGG pathways (**Table S2**). GO classification indicated comparable functional distributions across tissues, with the rhizome and flower containing 34,621 and 38,262 annotated transcripts, respectively (**Fig. 3A**). Roots and leaves exhibited similar GO profiles, suggesting a conserved functional repertoire (**Fig. S1**).

**Figure 3.**
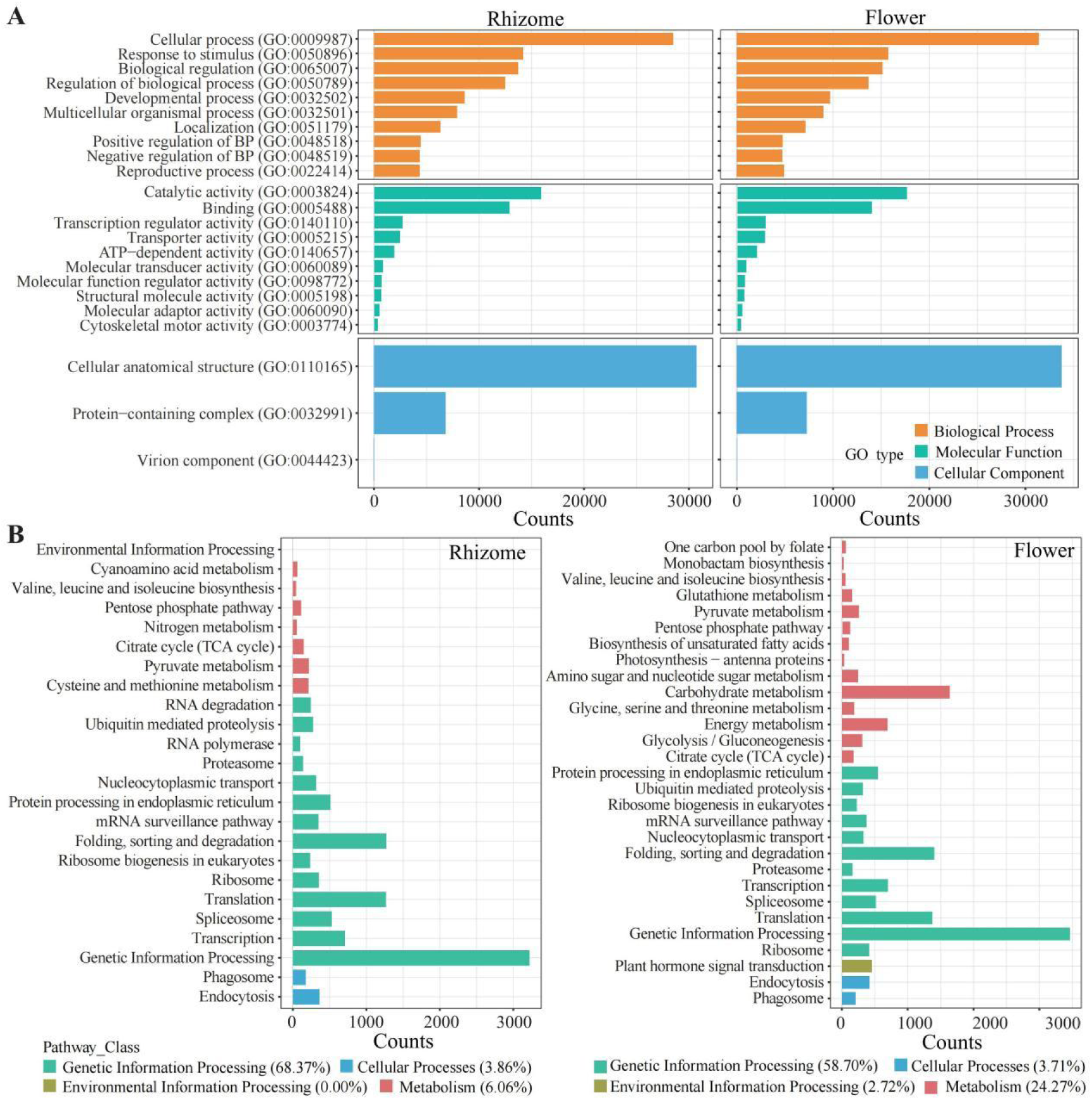
Functional annotation and classification of *P. vietnamensis* var. *fuscidiscus* transcriptomes in rhizome and flower. (A) Gene Ontology (GO) classification of transcriptomes from rhizome and flower. (B) Kyoto Encyclopedia of Genes and Genomes (KEGG) pathway classification of transcriptomes from rhizome and flower.

KEGG analysis showed that genetic information processing was the predominant functional category across all tissues. In contrast, the relative contribution of metabolism-related pathways varied among tissues, accounting for 24.27% in flower, 17.91% in leaf, 12.45% in root, 9.53% in stem, and 6.06% in rhizome (**Fig. 3B; Fig. S1**). Among metabolism-related subclasses, energy metabolism pathways were detected in flower (690 counts, 4.13%), leaf (640 counts, 4.34%), and stem (605 counts, 4.22%), but were not identified in root or rhizome. Together, these results suggest that while major functional classes are shared across tissues, the relative representation of metabolism-related pathways differs in a tissue-associated manner.

### Alternative splicing and implication in triterpene biosynthesis

A total of 56,019 expressed genes were identified, among which 45.68% produced two or more isoforms, indicating extensive alternative splicing (**Fig. 4A**). We detected 46,917 alternative splicing (AS) events, with retained intron (RI) being the most common type (44.51%), followed by alternative 3′ splice sites (A3, 19.61%), alternative 5′ splice sites (A5, 13.77%), and skipped exon (SE, 9.96%) (**Fig. 4B**, **4C**). Notably, key genes involved in triterpene biosynthesis exhibited considerable isoform diversity, suggesting that alternative splicing may contribute to the regulation of enzymatic activity. For example, *CYP716A53v2* displayed retained intron (RI) events (**Fig. 4D**), which could produce truncated or non-functional proteins and thereby modulate metabolic flux. Meanwhile, *dammarenediol-II synthase* (*DDS*) and *mevalonate diphosphate decarboxylase* (*MVD*) showed exon-skipping (ES) patterns (**Fig. S2A, B**), potentially generating isoforms with altered catalytic domains, which may be associated with tissue-dependent differences in triterpene biosynthesis. These observations suggest that alternative splicing may play a regulatory role in fine-tuning triterpene production across different tissues.

**Figure 4.**
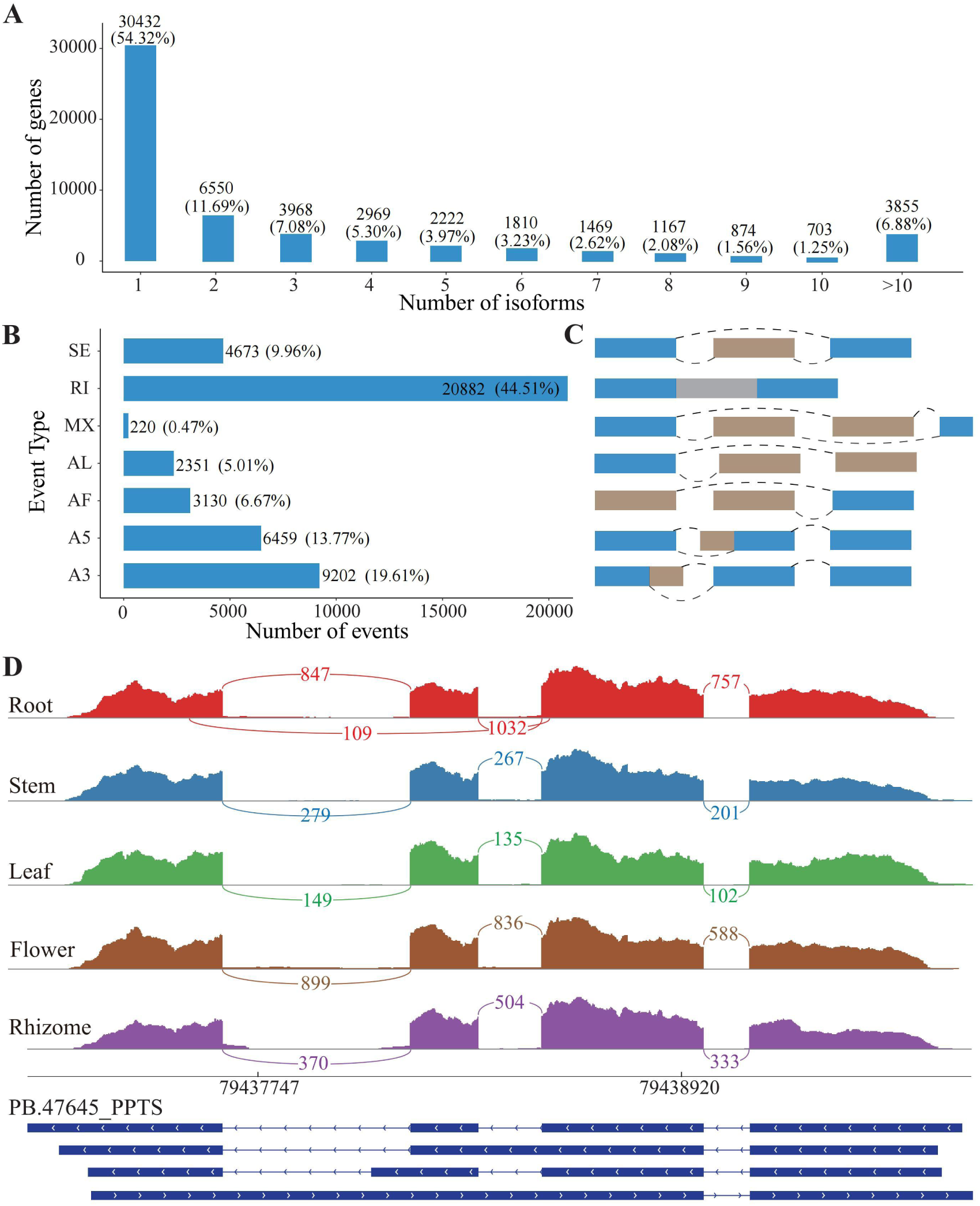
Characterization of alternative splicing (AS) events in *P. vietnamensis* var. *fuscidiscus*. (A) Proportion of genes producing one or more isoforms in the Iso-Seq data; counts and percentages are indicated above bars. (B) Total number of AS events identified from Iso-Seq data; counts and percentages are shown above bars. (C) Schematic representation of seven major AS types: skipped exon (SE), retained intron (RI), mutually exclusive exon (MX), alternative 3′ splice site (A3), alternative 5′ splice site (A5), alternative first exon (AF), and alternative last exon (AL). (D) Sashimi plot showing transcript isoforms of the cytochrome P450 gene *CYP716A53v2* (*PPTS*) based on Iso-Seq data. Peaks in red, blue, green, brown, and purple represent short-read RNA-seq coverage in root, stem, leaf, flower, and rhizome, respectively, and curved lines with numbers in corresponding colors represent splicing junctions supported by that number of short-read RNA-seq reads. For each isoform, blocks in blue represent exons and lines in-between represent introns. The presence of retained intron (RI) isoforms indicates alternative splicing–mediated transcript diversity in *CYP716A53v2*.

Further validation using RNA-Seq data across different tissues revealed substantial differential AS events (|ΔPSI| > 0.3) when comparing flower, leaf, root, and stem to rhizome. Specifically, 11,470, 11,323, 11,250, and 11,172 significant AS events were detected in flower, leaf, root, and stem compared to rhizome, respectively, each involving over 5,500 genes (**Fig. S2C**). RI and SE remained the predominant AS types across all comparisons (**Fig. S2D**), suggesting their potential regulatory roles in tissue-specific metabolic processes.

### Tissue-specific expression of genes involved in triterpene biosynthesis

To investigate tissue-specific gene expression, we analyzed transcriptomes of 281,468 unigenes and identified 14,427 differentially expressed genes (DEGs) across four pairwise comparisons with rhizome. The leaf *vs.* rhizome comparison had the highest number of DEGs (6,696), including 3,280 upregulated and 3,416 downregulated genes (**Fig. 5A**, **Fig. S3**). In contrast, the root vs. rhizome comparison showed the fewest DEGs (305), including 164 upregulated and 141 downregulated genes (**Fig. S3C**). Venn diagram analysis revealed distinct sets of DEGs across tissues, with a small core set shared across all comparisons (**Fig. 5B**).

**Figure 5.**
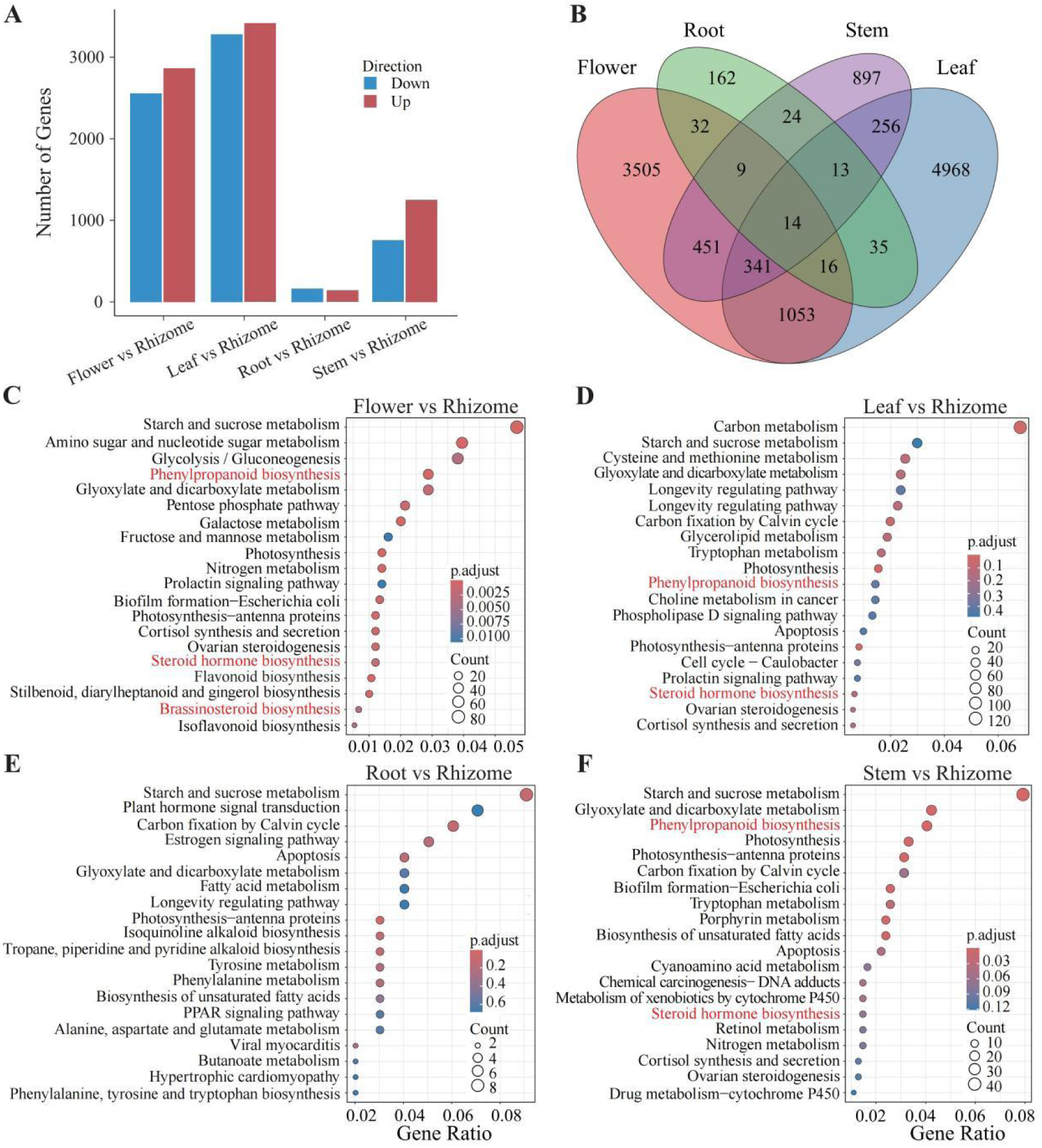
Tissue-specific differential gene expression and KEGG pathway enrichment in *P. vietnamensis* var. *fuscidiscus*. (A) Number of differentially expressed genes (DEGs) in pairwise comparisons between rhizome and other tissues; red and blue bars represent up- and down-regulated genes, respectively. (B) Venn diagram showing overlap of DEGs across four tissue comparisons (flower, leaf, root, and stem vs. rhizome). (C–F) Top 20 KEGG pathways enriched in DEGs for flower vs. rhizome (C), leaf vs. rhizome (D), root vs. rhizome (E), and stem vs. rhizome (F). Pathways related to phenylpropanoid biosynthesis and steroid hormone biosynthesis are highlighted in red.

KEGG enrichment analysis of DEGs from comparisons between each tissue and the rhizome identified the top 20 significantly enriched KEGG pathways, suggesting pronounced tissue-associated differences in metabolic pathway representation (**Fig. 5C-F**). Pathways related to plant defense and triterpenoid precursor biosynthesis, including phenylpropanoid biosynthesis and steroid hormone biosynthesis, were significantly enriched in flower, leaf, and stem compared with the rhizome, whereas no significant enrichment was observed in the root versus rhizome comparison.

Because KEGG enrichment reflects pathway-level overrepresentation rather than gene expression levels, we further examined the expression patterns of genes involved in these two pathways. Genes related to phenylpropanoid biosynthesis displayed diverse expression patterns across tissues and did not show preferentially high expression in rhizome or root (**Fig. S4A**). In contrast, steroid hormone biosynthesis–related genes exhibited generally higher expression levels in rhizome and root, with lower expression in flower, leaf, and stem (**Fig. S4B**). As this pathway produces 2,3-oxidosqualene, a key precursor for triterpenoid saponin biosynthesis, these results are consistent with the rhizome and root being important tissues associated with saponin biosynthesis and accumulation in *P. vietnamensis*.

### Expression patterns of triterpene saponin biosynthetic genes

To further characterize triterpenoid biosynthesis, we identified 21 enzyme-encoding genes involved in the pathway, covering the upstream mevalonate (MVA) pathway, the branching step from 2,3-oxidosqualene, and the downstream oleanane (OA)-, protopanaxatriol (PPT)-, ocotillol (OCT)-, and protopanaxadiol (PPD)-type routes (**Fig. 6A**; **Table S3**).

**Figure 6.**
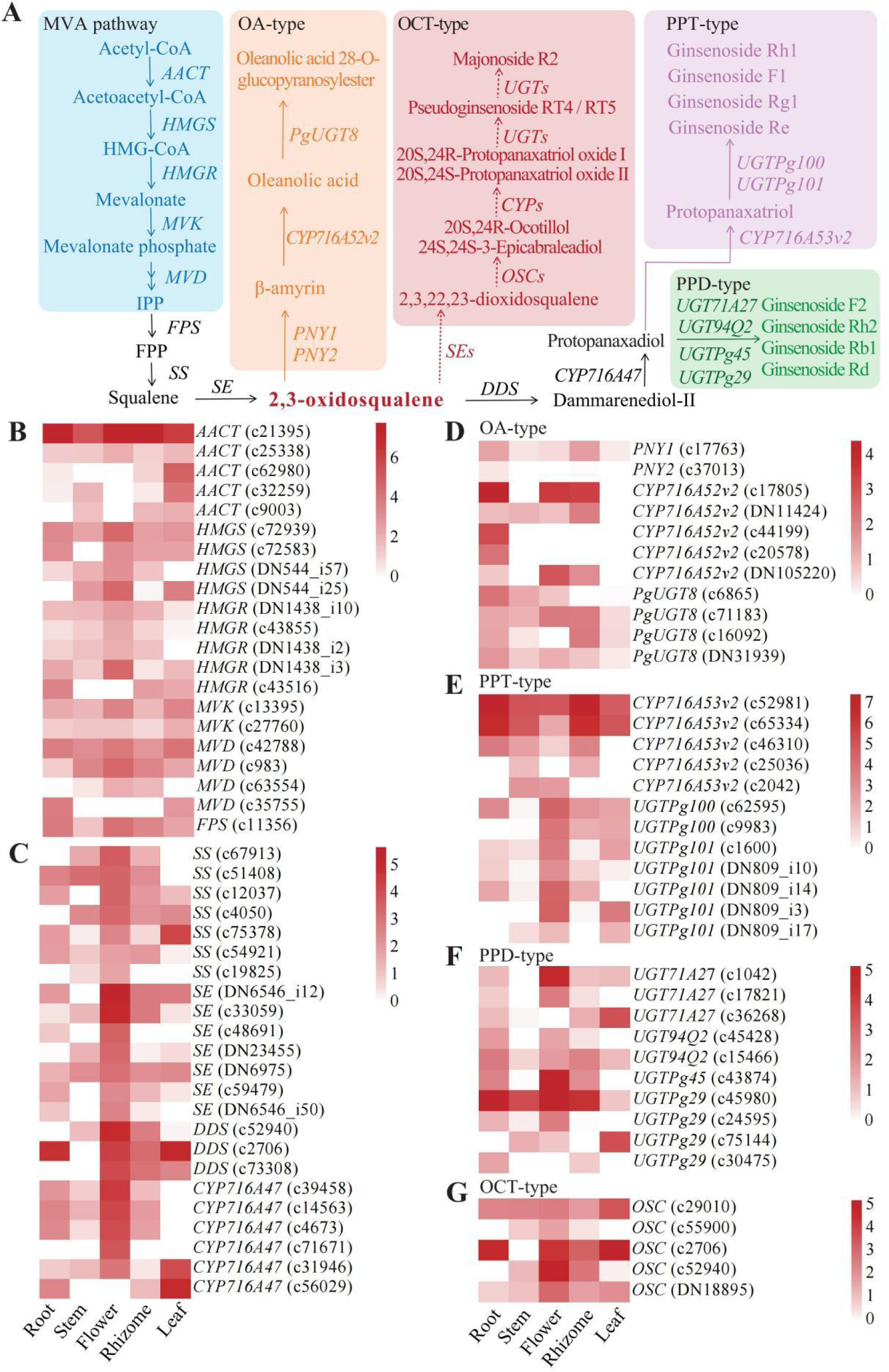
Triterpenoid saponin biosynthesis pathways and tissue-specific expression of candidate genes in *P. vietnamensis* var. *fuscidiscus*. (A) Proposed biosynthetic pathways from the mevalonate (MVA) pathway to oleanane (OA)-type, protopanaxadiol (PPD)-type, protopanaxatriol (PPT)-type, and ocotillol (OCT)-type saponins. Candidate genes are shown in italics. (B–F) Heatmaps showing tissue-specific expression patterns of candidate genes involved in different steps of triterpenoid saponin biosynthesis, including enzymes from the upstream MVA pathway and homologous genes associated with the biosynthetic pathways of OA-type, PPD-type, PPT-type, and OCT-type saponins. (G) Heatmap showing the expression patterns of all identified oxidosqualene cyclase (*OSC*) genes across five tissues (root, stem, flower, leaf, and rhizome). Gene expression levels are shown as log_2_(TPM + 1). The abbreviations are indicated as follows, *AACT*: acetyl-CoA acetyltransferase, *HMGS*: 3-hydroxy-3-methylglutaryl coenzyme A synthase, *HMGR*: 3-hydroxy-3-methylglutaryl coenzyme A reductase, *MVK*: mevalonate kinase, *MVD*: mevalonate diphosphate decarboxylase, *FPS*: farnesyl diphosphate synthase, *SS*: squalene synthase, *SE*: squalene epoxidase, *DDS*: dammarenediol synthase, *CYP*: cytochrome P450, *UGT*: UDP-glycosyltransferase.

Expression analysis across tissues revealed distinct patterns (**Fig. 6B–F**). Genes of the MVA pathway (e.g., *AACT*, *HMGS*, *HMGR*, *MVK*, *MVD*) were broadly expressed, while *FPS*, *SS*, *SE*, *DDS*, and *CYP716A52* showed highest expression in flowers, suggesting relatively higher precursor biosynthetic activity in this organ (**Fig. 6B, 6C**). Within the OA-type branch, β-amyrin synthases (*PNY1*, *PNY2*) and *CYP716A52v2* were predominantly expressed in roots, suggesting a root-associated preference in OA-type saponin biosynthesis. Similarly, the UDP-glycosyltransferase *PgUGT8* was more highly expressed in roots (**Fig. 6D**).

In contrast, PPT-type biosynthetic genes such as *CYP716A53v2*, *UGTPg100*, and *UGTPg101* exhibited flower-specific expression, consistent with the DEG enrichment observed in the flower vs. rhizome comparison (**Fig. 5C**). This suggests that hydroxylation and glycosylation steps in PPT saponin formation occur mainly in floral tissues (**Fig. 6E**). Meanwhile, PPD-type glycosyltransferases including *UGT71A27*, *UGT94Q2*, *UGTPg45*, and *UGTPg29* did not exhibit strong tissue specificity, indicating that PPD-type glycosylation may occur broadly in both aerial and underground organs (**Fig. 6F**).

In addition to these classical branches, we further examined the expression profiles of all candidate *oxidosqualene cyclases* (*OSCs*) as well as *UGTs* potentially involved in the terminal modification steps of OCT-type saponins such as majonoside R2. All five *OSC* candidates were detectable across tissues, with slightly lower expression in stems but no clear organ-enriched pattern overall (**Fig. 6G**). Members of the *UGT* gene family also showed dispersed and heterogeneous expression across the five tissues, and no consistent co-expression pattern with the *OSC* candidates was observed (**Fig. S5**). These results suggest that, although these *OSCs* and *UGTs* represent plausible candidates for the final cyclization and glycosylation steps of OCT-type saponin biosynthesis, their transcriptional profiles do not indicate strong organ specialization. This implies that these modifications may occur in multiple tissues or be influenced by post-transcriptional or enzymatic regulatory mechanisms.

Overall, triterpene saponin biosynthesis in *P. vietnamensis* exhibits apparent tissue-associated patterns of pathway activity: OA-type saponins are primarily synthesized in roots, PPT-type modifications occur mainly in flowers, and PPD-type glycosylation takes place across multiple tissues. In contrast, candidate genes potentially involved in OCT-type saponin biosynthesis exhibit broad and non-specific expression patterns, suggesting that the terminal tailoring steps of this characteristic saponin class may occur in various organs.

## Discussion

In this study, we integrated long-read PacBio Iso-Seq and short-read Illumina RNA-Seq to construct the first high-quality, full-length transcriptome atlas of *P. vietnamensis* var. *fuscidiscus* across five tissues. This comprehensive dataset uncovered widespread alternative splicing, extensive transcript diversity, and robust expression of genes involved in ginsenoside biosynthesis throughout the five major tissues. In contrast to a previous Illumina-based transcriptome study that assembled 126,758 unigenes with an average length of 1,304 bp and no isoform information [19], our full-length transcriptome identified 281,468 non-redundant transcripts at isoform resolution, including 8,089 novel genes and 46,917 alternative splicing events. This represents a nearly 2.2-fold increase in transcript number and demonstrates the capability of long-read sequencing in revealing intricate isoform diversity, providing new insights into triterpenoid biosynthesis. Furthermore, we identified 21 key enzyme-encoding genes potentially involved in the biosynthesis of oleanane-type (OA), protopanaxadiol-type (PPD), protopanaxatriol-type (PPT), and ocotillol-type (OCT) triterpenoid saponins. These findings offer important molecular insights into the exceptional accumulation of OCT ginsenosides—particularly majonoside R₂—in this species [1, 3].

### Tissue-specific specialization of triterpenoid biosynthetic pathways

Our transcriptomic analysis revealed that upstream mevalonate (MVA) pathway genes—including *AACT*, *HMGR*, *MVK*, and *MVD*—were expressed constitutively in all tissues, consistent with their essential role in supplying precursors for triterpenoid biosynthesis. In contrast, downstream branch-specific genes exhibited distinct tissue-specific expression patterns. The enriched expression of β-amyrin synthases (*PNY1*, *PNY2*) and *CYP716A52v2* in roots suggests active biosynthesis of OA-type saponins in underground tissues. Conversely, *CYP716A53v2* and glycosyltransferases such as *UGTPg100* and *UGTPg101* were predominantly expressed in flowers, suggesting that hydroxylation and glycosylation steps for PPT-type saponins may preferentially occur in floral tissues. PPD-type glycosyltransferases (e.g., *UGT94Q2*, *UGT71A27*, *UGTPg29*) were more broadly distributed across multiple tissues, implying widespread PPD saponin production. These expression patterns are consistent with previously reported metabolite profiles [54], further supporting the tissue-specific compartmentalization of triterpenoid biosynthesis in *P. vietnamensis*.

### Alternative splicing and functional diversification

The integration of Iso-Seq and RNA-Seq data revealed a high prevalence of alternative splicing (AS), with more than 45% of expressed genes producing multiple isoforms. Retained intron (RI) emerged as the most common type of AS event. Notably, key genes including *CYP716A53v2*, *DDS*, and *MVD* exhibited tissue-specific splicing patterns. These AS events may influence enzyme function, subcellular localization, or stability, thereby contributing to the precise regulation of triterpene biosynthesis. For instance, RI-type splicing in *DDS* could generate distinct protein isoforms with altered catalytic properties, potentially influencing ocotillol scaffold formation. These results highlight the role of post-transcriptional regulation in the sophisticated control of specialized metabolic pathways.

The widespread alternative splicing observed in *P. vietnamensis* var. *fuscidiscus* (48% multi-isoform genes) is consistent with long-read transcriptome studies across diverse species. In closely related Panax species, SMRT sequencing of P. notoginseng uncovered 20,015 AS events from 6,324 genes and 8,111 novel isoforms [27], and multi-tissue Iso-Seq of P. ginseng similarly revealed extensive isoform expansion [55]. Comparable patterns are reported in other dicots: 93.1% multi-isoform genes in *Camellia oleifera* seed [56], 40.6–52.8% in *Phragmites australis* [34], 59.5% in *Brassica napus* with an average of 3.81 isoforms per locus [57], 34.5–36.5% in *Astragalus membranaceus* [58], and 50.83% in *Gossypium australe* [59]. Animal long-read transcriptomes also show similar complexity—for example, 3.91 isoforms per gene (60.93% multi-isoform loci) in rat hippocampus [32], and 25–65% multi-isoform genes in threespine stickleback fish [33] and pig testis cells [60]. Collectively, long-read transcriptomes typically reveal 30–60% of genes with multiple isoforms and an average of 2–4 isoforms per locus. Therefore, the isoform diversity we detected in *P. vietnamensis* falls within the range reported for both *Panax* and other dicot plants, reinforcing that alternative splicing is a major driver of transcriptomic and functional diversification in this lineage.

### Mechanistic implications for OCT-type ginsenoside accumulation

Despite the pharmacological significance of ocotillol (OCT)-type ginsenosides, particularly their anticancer and hepatoprotective properties, the biosynthetic pathway responsible for formation of the OCT scaffold remains poorly understood. [17]. The distinctive structure of these saponins, characterized by a tetrahydrofuran (THF) ring, suggests the involvement of an atypical *oxidosqualene cyclase* (*OSC*) that differs from canonical β-amyrin synthase or dammarenediol synthase. Although no *OSC* specific to OCT biosynthesis was identified among the annotated transcripts, the detection of alternatively spliced variants of *DDS* may point to possible neofunctionalization. Further biochemical validation using heterologous expression systems will be essential to identify the *OSC* that catalyzes the cyclization of 2,3-oxidosqualene into the OCT backbone.

We further investigated the biosynthesis of majonoside R2, the hallmark OCT-type saponin of *P. vietnamensis*. Prior biochemical studies demonstrated that *PvfUGT1* and *PvfUGT2* sequentially catalyze the two glycosylation steps converting protopanaxatriol-oxide intermediates (RT4/RT5) into majonoside R2, providing key enzymatic evidence for this pathway [61, 62]. To place these enzymes within the broader *UGT* repertoire of *P. vietnamensis*, we incorporated *PvfUGT1* and *PvfUGT2* into a phylogenetic framework and examined the expression of candidate *UGTs* clustering alongside them. These *UGTs* showed broad, non-specialized expression across all tissues, and the same pattern was observed for candidate *OSCs* that may catalyze OCT backbone formation. The absence of tissue enrichment suggests that majonoside R2 biosynthesis is not confined to a specific organ and may depend on additional regulatory factors, such as substrate availability and subcellular compartmentation, rather than on transcript abundance alone.

Together, these findings indicate that, unlike the clearly partitioned OA-, PPT-, and PPD-type pathways, the OCT-type pathway is more diffusely regulated. Integrating established biochemical functions of *PvfUGT1* and *PvfUGT2* with our phylogenetic and expression analyses refines the current model of MR2 biosynthesis and highlights targets for future functional and metabolic engineering studies.

### Evolutionary adaptations and biosynthetic innovation

From an evolutionary perspective, the ability of *P. vietnamensis* to accumulate up to 75% ocotillol (OCT)-type saponins in its rhizomes likely arises from multiple synergistic mechanisms: (1) constitutive expression of upstream MVA pathway genes in underground tissues ensures a steady supply of precursors; (2) neofunctionalized oxidosqualene cyclase (OSC) enzymes or alternatively spliced *DDS* isoforms may enable ocotillol production; and (3) an extensive glycosyltransferase network facilitates diverse modifications of the ocotillol scaffold. These adaptations collectively suggest that OCT biosynthesis in *P. vietnamensis* may represent a lineage-specific metabolic innovation, potentially driven by gene duplication and functional divergence following whole-genome duplication (WGD), as proposed in other *Panax* species.

While this study provides a comprehensive transcriptomic resource for *P. vietnamensis* var. *fuscidiscus* and identifies candidate genes and alternative splicing events potentially involved in OCT-type ginsenoside biosynthesis, functional validation of these genes (e.g., knock-down, knock-out, overexpression) was not performed. Key knowledge gaps remain, including the identity of the *OSCs* responsible for forming the OCT skeleton. Nevertheless, the high-quality transcriptome and tissue-specific expression profiles generated here provide a solid foundation for targeted functional studies. Future work should focus on gene characterization via CRISPR/Cas-mediated knockout, overexpression, or heterologous expression, and on elucidating transcriptional regulators and cis-elements controlling tissue-specific expression. Integrating transcriptomic, metabolomic, proteomic, and single-cell RNA sequencing data will further enable a systems-level understanding of OCT biosynthesis and reveal new targets for metabolic engineering.

In conclusion, our study provides a comprehensive transcriptomic resource for *P. vietnamensis* and marks a substantial step toward deciphering the biosynthetic pathways and regulatory mechanisms underlying OCT ginsenoside accumulation. This work not only enhances our understanding of ginseng metabolic diversity, but also establishes a foundation for synthetic biology approaches aimed at the sustainable production of pharmacologically valuable OCT saponins.

## Conclusion

This study reports the first high-quality, full-length, tissue-specific transcriptome atlas of *P. vietnamensis* var. *fuscidiscus*, constructed through integrated PacBio Iso-Seq and Illumina RNA-Seq sequencing. A total of 281,468 non-redundant transcripts were assembled, including 8,089 novel genes not previously annotated, substantially expanding the currently available genetic repertoire of this pharmacologically valuable species.

We identified 21 key enzyme-encoding genes involved in triterpenoid saponin biosynthesis and identified tissue-associated expression patterns across the oleanane-, protopanaxadiol-, protopanaxatriol-, and ocotillol-type biosynthetic pathways. Notably, over 46,000 alternative splicing events were detected, including isoform diversity in critical biosynthetic genes such as *DDS* and *CYP716A53v2*, underscoring the importance of post-transcriptional regulation in specialized metabolite production. Collectively, these findings provide new molecular insights relevant to ocotillol-type ginsenoside biosynthesis and establish a valuable transcriptomic resource that may support future studies aimed at identifying candidate enzymes, exploring potential regulatory mechanisms, and guiding subsequent functional and metabolic engineering research. This comprehensive transcriptomic resource also reinforces the value of *P. vietnamensis* as an excellent model for studying lineage-specific metabolic innovations in medicinal plants.

## Abbreviations

AACT: Acetyl-CoA acetyltransferase
AF: Alternative first exon
AL: Alternative last exon
AS: Alternative splicing
A3: Alternative 3′ splice site
A5: Alternative 5′ splice site
CCS: Circular consensus sequence
CD-HIT: Cluster Database at High Identity with Tolerance
COG/KOG: Clusters of Orthologous Groups / Eukaryotic Orthologous Groups
CYP: Cytochrome P450
DDS: Dammarenediol synthase
DEG: Differentially expressed gene
eggNOG: Evolutionary genealogy of genes: Non-supervised Orthologous Groups
FDR: False discovery rate
FL: Full-length
FLNC: Full-length non-chimeric
FPS: Farnesyl diphosphate synthase
FSM: Full splice match
GO: Gene Ontology
HMGR: 3-hydroxy-3-methylglutaryl-coenzyme A reductase
HMGS: 3-hydroxy-3-methylglutaryl coenzyme A synthase
HQ: High-quality
ICE: Iterative Clustering for Error Correction
ISM: Incomplete Splice Match
Iso-Seq: Isoform sequencing
KEGG: Kyoto Encyclopedia of Genes and Genomes
LQ: Low-quality
MVA: Mevalonate pathway
MVD: Mevalonate diphosphate decarboxylase
MVK: Mevalonate kinase
MX: Mutually exclusive exons
NIC: Novel in catalog
NNC: Novel not in catalog OCT Ocotillol
OSC: Oxidosqualene cyclase
PPD: Protopanaxadiol
PPT: Protopanaxatriol
RI: Retained intron
RNA-Seq: RNA sequencing
ROIs: Reads of insert
SE: Skipped exon
SE: Squalene epoxidase
SGS: Second-generation sequencing
SMRT: Single-Molecule Real-Time sequencing
SS: Squalene synthase
SSR: Simple sequence repeat
TSS/TTS: Transcription start site / transcription termination site
UGT: UDP-glycosyltransferase

## Authors’ contributions

LZG conceived the study. WL performed transcriptome analyses and drafted the manuscript. ZX and AN contributed to RNA extraction, sequencing, and data validation. LZG revised the manuscript critically and supervised the project. All authors read and approved the final manuscript.

## Acknowledgements

Not applicable.

## Data availability

The raw data were submitted to NCBI under the BioProject number PRJNA1312250.

## Declarations

### Ethics approval and consent to participate

Not applicable.

### Consent for publication

Not applicable.

### Clinical trial number

Not applicable.

## Supplementary Information

**Figure S1. Functional annotation and classification of *P. vietnamensis* var. *fuscidiscus* transcriptomes in leaf, root, and stem.** (A, B) GO terms and KEGG pathways assigned to leaf transcriptomes. (C, D) GO terms and KEGG pathways assigned to root transcriptomes. (E, F) GO terms and KEGG pathways assigned to stem transcriptomes.

**Figure S2. Comparative analysis of alternative splicing (AS) events across tissues in *P. vietnamensis* var. *fuscidiscus*.** (A) Sashimi plot illustrating isoform structures of the *dammarenediol synthase* (*DDS*) gene based on Iso-Seq data. (B) Sashimi plot illustrating isoform structures of the *mevalonate-5-pyrophosphate decarboxylase* (*MVD*) gene based on Iso-Seq data. In panels (A) and (B), rectangular boxes represent exons, with box lengths proportional to exon size, while connecting lines indicate splicing junctions. Numbers displayed on the junction arcs denote the number of full-length Iso-Seq reads supporting each splicing event. Colored coverage peaks (red, blue, green, brown, and purple) represent short-read RNA-seq coverage in root, stem, leaf, flower, and rhizome, respectively. Curved lines with numbers in corresponding colors indicate splice junctions supported by short-read RNA-seq data in each tissue. For each isoform, blue blocks represent exons and the connecting lines indicate introns. (C) Counts of significant AS events (|ΔPSI| > 0.3) and associated genes in pairwise comparisons between flower, leaf, root, or stem and rhizome. (D) Distribution of seven major AS event types (SE, RI, MX, A5, A3, AF, AL) across tissue comparisons. RI and SE were the most abundant types.

**Figure S3. MA plots of differential gene expression between rhizome and other tissues in *P. vietnamensis* var. *fuscidiscus*.** (A–D) MA plots for flower (A), leaf (B), root (C), and stem (D) versus rhizome. Log₂ fold change (logFC) is plotted against average expression (logCounts). Significantly differentially expressed genes (DEGs) are shown in red; non-significant genes are in black.

**Figure S4. Expression patterns of genes involved in phenylpropanoid and steroid hormone biosynthesis across tissues.** (A) Heatmap showing the expression patterns of genes involved in phenylpropanoid biosynthesis (KEGG pathway 00940) across five tissues (root, stem, flower, leaf, and rhizome). (B) Heatmap showing the expression patterns of genes involved in steroid hormone biosynthesis (KEGG pathway 00140) across five tissues. Gene expression levels are shown as log₂(TPM + 1). Genes are displayed without hierarchical clustering to facilitate direct comparison of expression levels among tissues.

**Figure S5. Expression patterns of *UDP-glycosyltransferase* (*UGT*) genes phylogenetically related to *PvfUGT1* and *PvfUGT2* in *P. vietnamensis* var. *fuscidiscus*.** Heatmap showing the expression profiles of *UGT* genes that cluster with the functionally characterized enzymes *PvfUGT1* and *PvfUGT2* in phylogenetic analysis. *PvfUGT1* and *PvfUGT2*, which have been previously demonstrated to catalyze the two sequential glycosylation steps leading to Majonoside R2 biosynthesis, are included as reference genes. Expression patterns of the phylogenetically related *UGTs* are shown across five tissues (root, stem, flower, leaf, and rhizome). Gene expression levels are presented as log₂(TPM + 1).

**Table S1. Summary of final reference transcripts assembled from integrated RNA-Seq and Iso-Seq data.**

**Table S2. Functional annotation statistics of unigenes across tissues and transcriptomes based on EggNOG, GO, COG/KOG, and KEGG databases.**

**Table S3. Candidate genes involved in triterpenoid saponin biosynthesis in *P. vietnamensis* var. *fuscidiscus*, including protein annotations, gene names, and transcript abundance (TPM) across root, stem, flower, rhizome, and leaf.**

