## Supplementary figures and images for "Full-length transcriptome atlas of *Panax vietnamensis* var. *fuscidiscus* reveals novel genes and alternative splicing in tissue-specific biosynthesis of ocotillol-type saponins"

### Figure S1

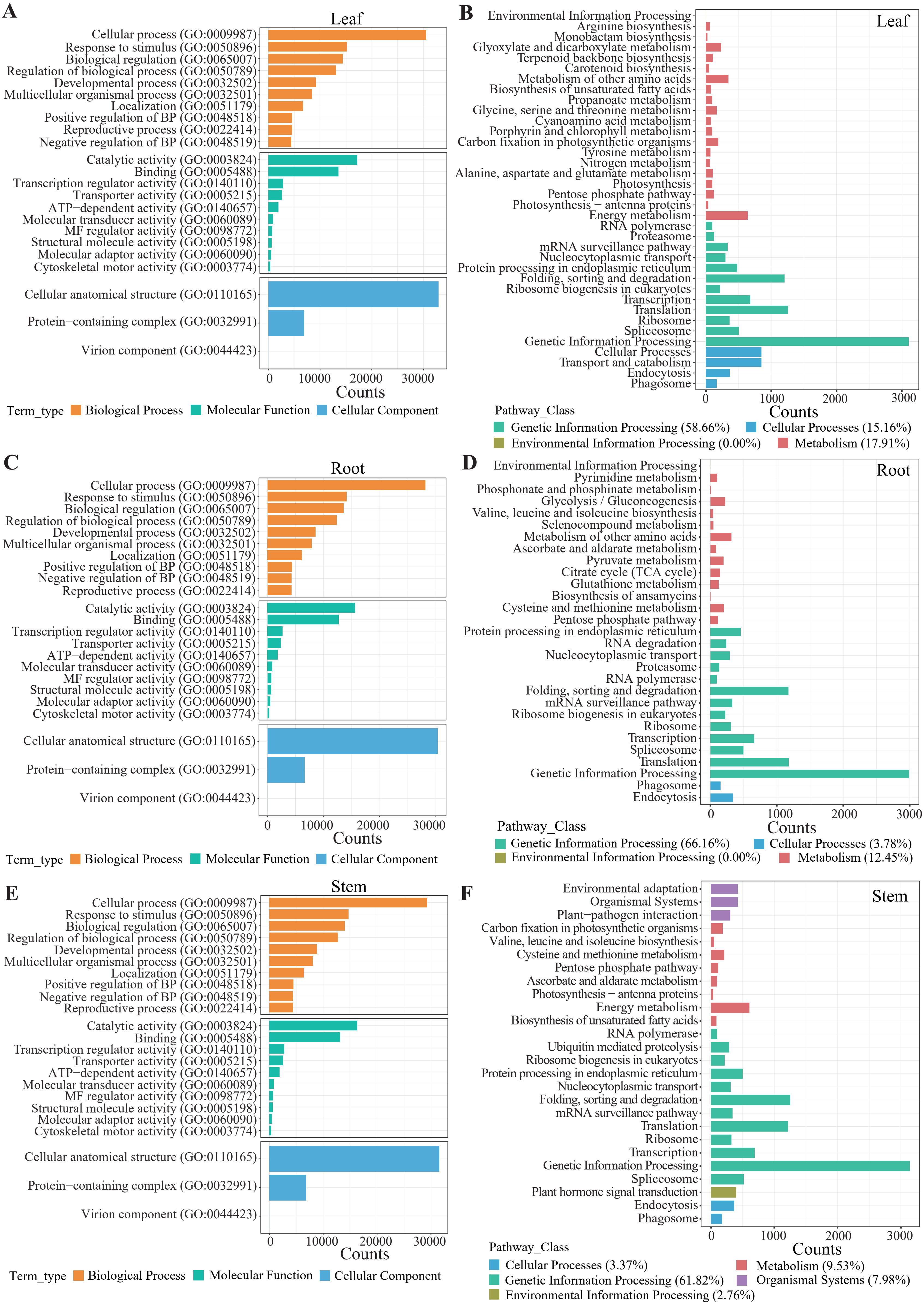

### Figure S2

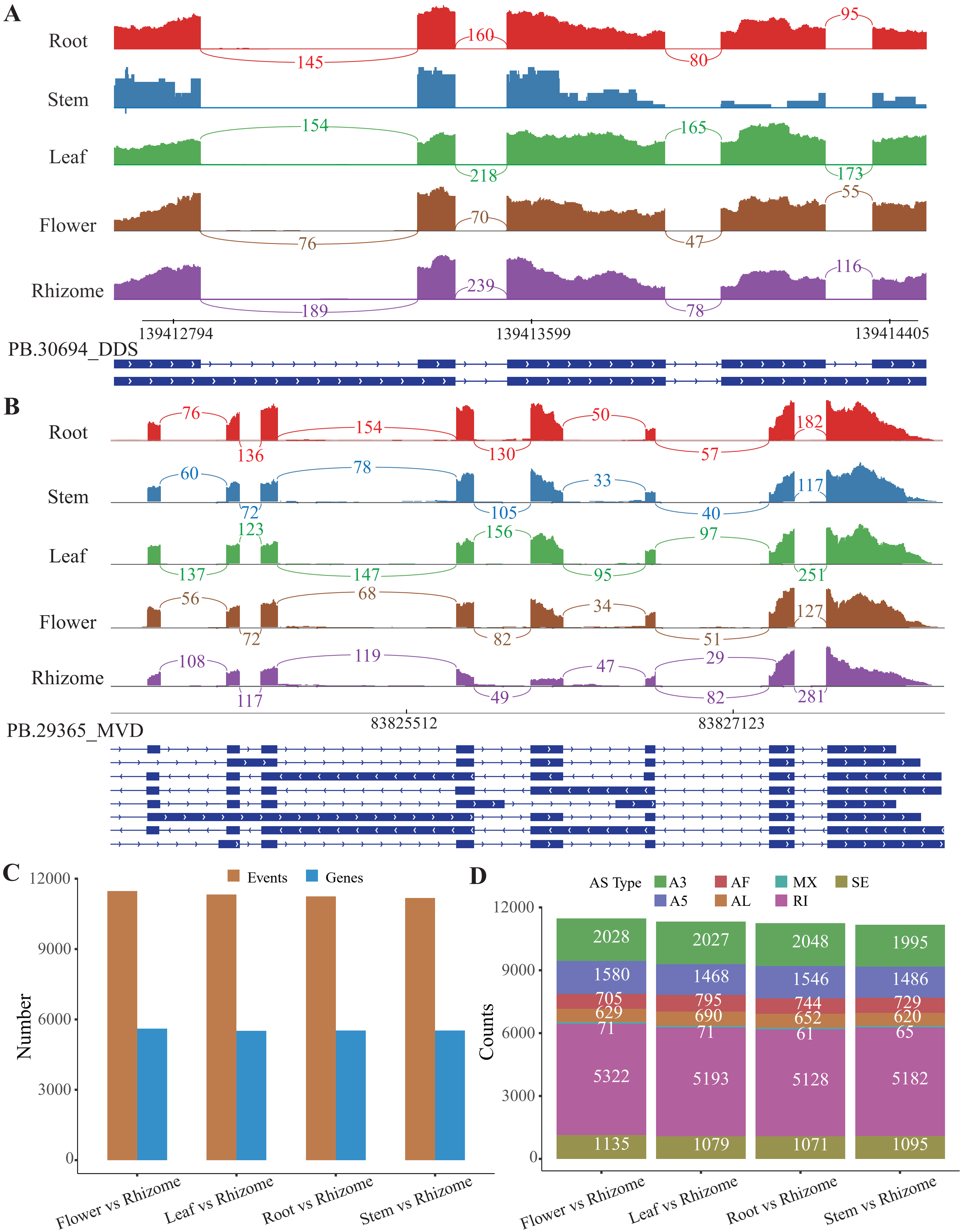

### Figure S3

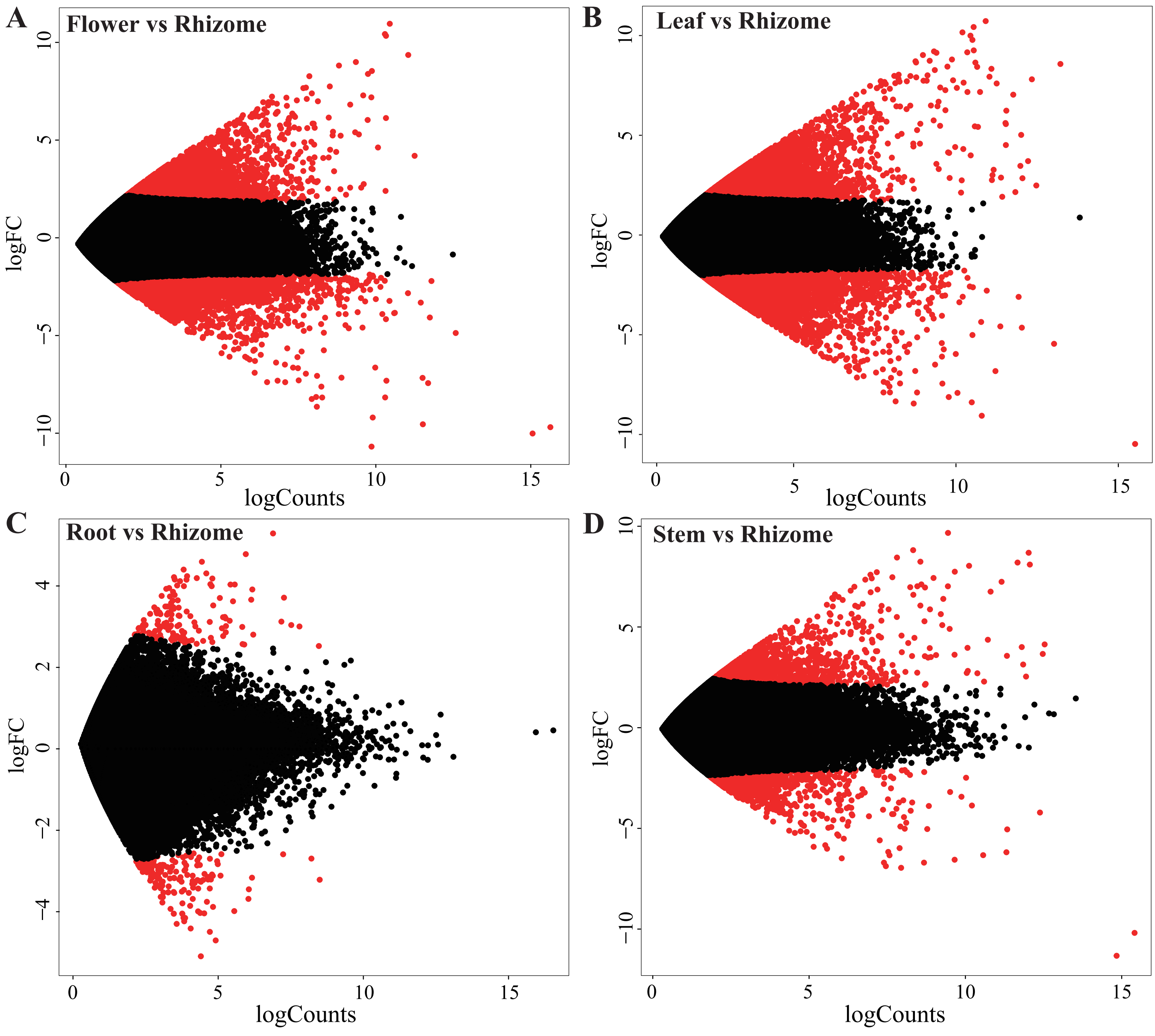

### Figure S4

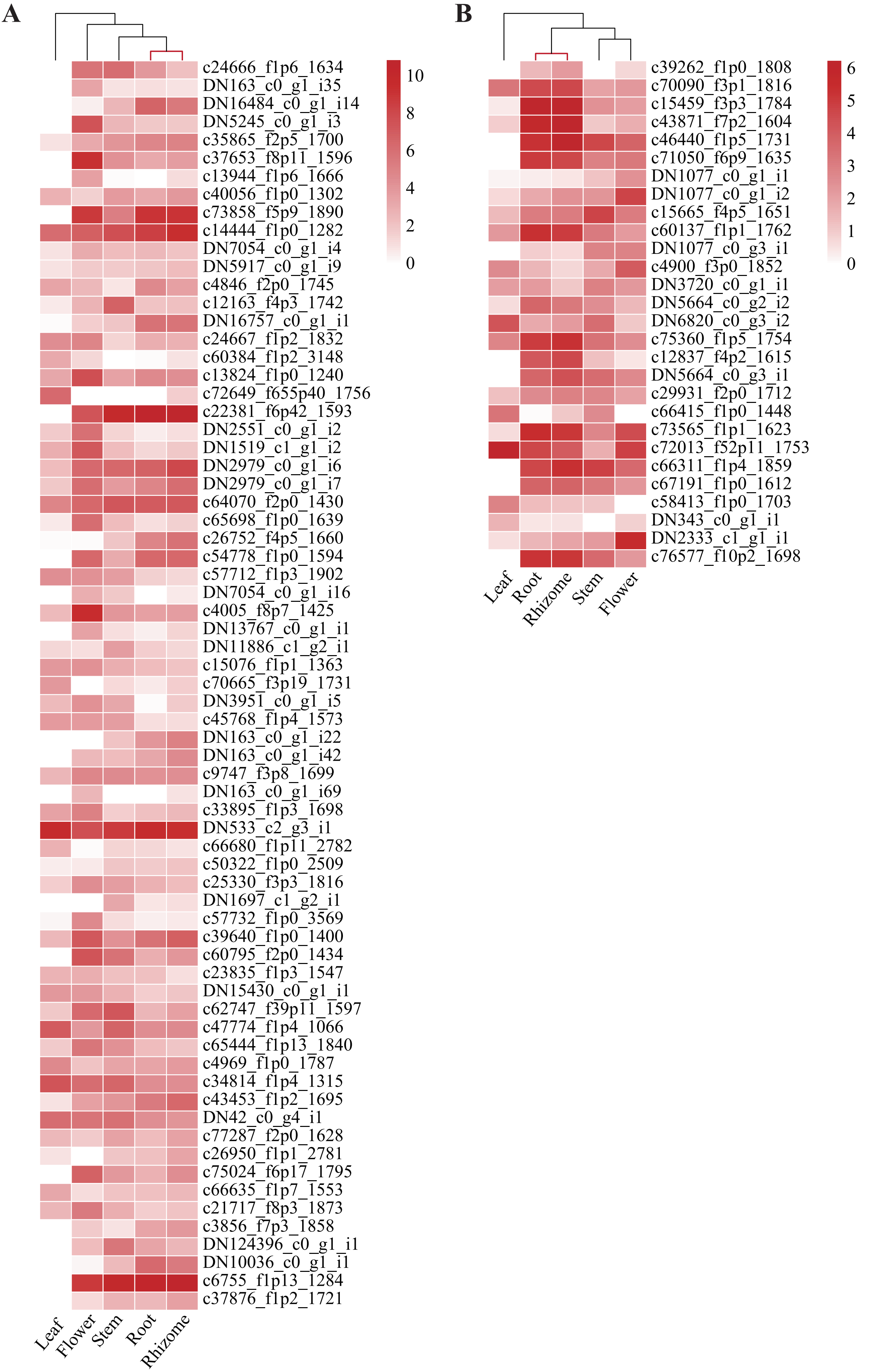

### Figure S5

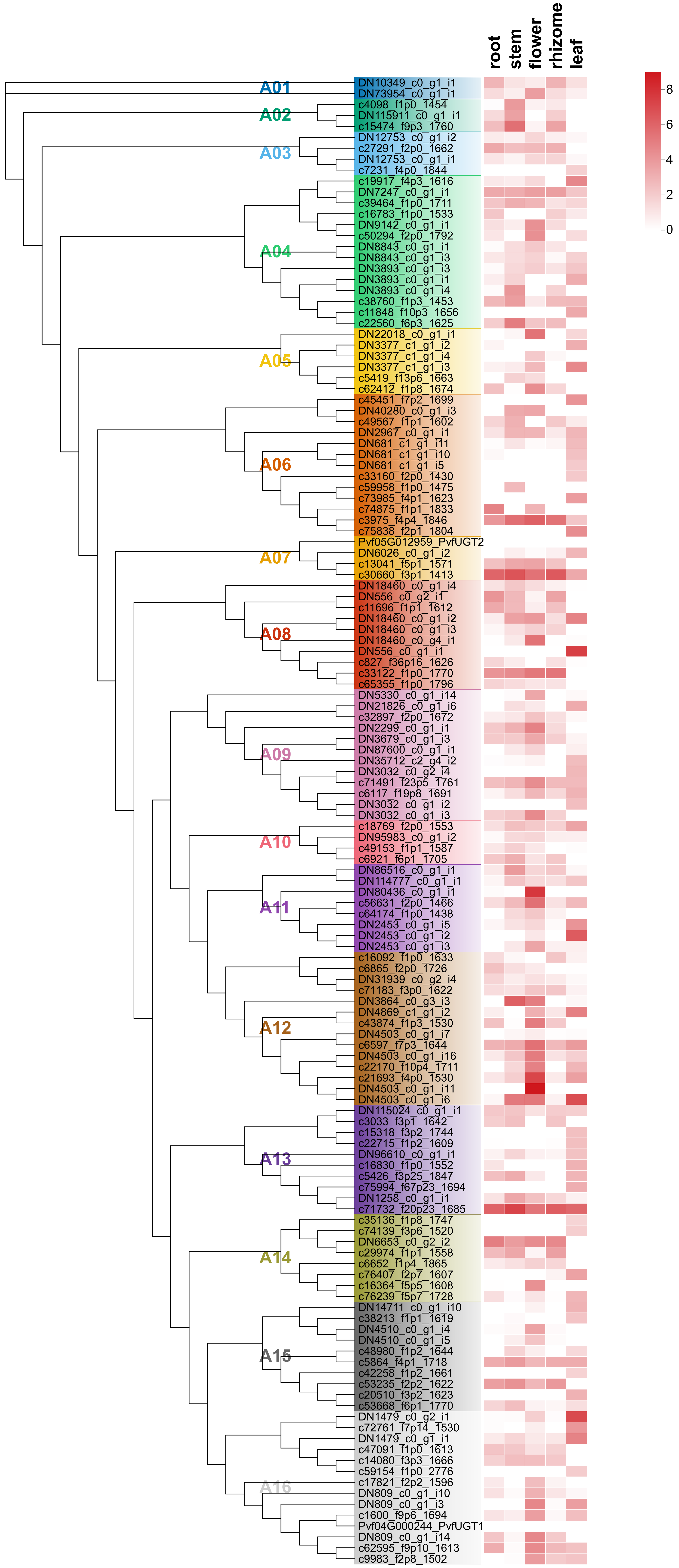
